# First genome-wide association study of non-severe malaria in two birth cohorts in Benin

**DOI:** 10.1101/483859

**Authors:** Jacqueline Milet, Anne Boland, Pierre Luisi, Audrey Sabbagh, Ibrahim Sadissou, Paulin Sonon, Nadia Domingo, Friso Palstra, Laure Gineau, David Courtin, Achille Massougbodji, André Garcia, Jean-François Deleuze, Hervé Perdry

## Abstract

Recent research efforts to identify genes involved in malaria susceptibility using genome-wide approaches have focused on severe malaria. Here we present the first GWAS on non-severe malaria designed to identify genetic variants involved in innate immunity or innate resistance mechanisms. Our study was performed on two cohorts of infants from southern Benin (525 and 250 individuals respectively) closely followed from birth to 18-24 months of age, with an assessment of a space-and time-dependent environmental risk of exposure. Both the recurrence of mild malaria attacks and the recurrence of malaria infections as a whole (symptomatic and asymptomatic) were considered. Our study highlights a role of *PTPRT*, a tyrosine phosphatase receptor involved in STAT3 pathway and several other genes whose biological functions are relevant in malaria infection. Results shows that GWAS on non-severe malaria can successfully identify new candidate genes and inform physiological mechanisms underlying natural protection against malaria.

## Introduction

In spite of numerous prevention and control efforts in recent years, malaria remains a major global public health problem with 219 million cases and ∼ 435,000 deaths in 2017 *(1)*. In Africa, Bhatt et *al. (2)* have estimated that *Plasmodium falciparum* infection prevalence halved between 2000 and 2015, and that the incidence of clinical disease fell by 40%, owing mainly to the large distribution of insecticide-treated nets, the most widespread intervention. However the fight against malaria is currently facing numerous challenges. A decrease in the global reduction of malaria cases and deaths is observed at present, with no significant progress made over the 2015–2017 period. The WHO African region which represents the region with the highest malaria burden (92% of malaria cases and 93% of malaria-related deaths in 2017) is also the area where medical advances have been hardest to achieve recently. Moreover control efforts are threatened by insecticide resistance and the possible spread of resistance to artemisinin (an essential component of the most efficient anti-malaria drugs, the artemisinin-based combination therapy-ACT) from Southeast Asia to Africa. In absence of alternative treatments with the same efficacy and tolerability as ACT and of an efficient vaccine, fundamental malaria research continues to be essential in order to better understand the physiopathology of the disease.

*P. falciparum* is the most prevalent malaria parasite in sub-Saharan Africa (99.7% of malaria cases in African region). Different clinical presentations of *P. falciparum* malaria exist, from asymptomatic (parasite carriage without any clinical sign), or mild forms (parasitemia with fever), to severe forms which may ultimately lead to death. This variability of clinical presentation is thought to be attributable to environmental factors, as well as parasite and human host factors, among which genetic variation could play a major role (3,4). In 2005, Mackinnon et al. (5) estimated that 24% and 25% of the variability in the incidence of mild malaria and severe malaria, respectively, could be explained by genetic factors. Numerous genetic epidemiological studies have attempted to identify gene polymorphisms associated with susceptibility or resistance to different malaria phenotypes (see (4,6,7)).

Most studies performed to date have focused on severe malaria, despite the fact that mild forms represent the major part of the global burden. Candidate gene studies for mild forms focused on genetic polymorphisms involved in host immune response (*IL10, IL3, LTA*,), in genes possibly involved in oxidative stress (*HP*), red blood cell (RBC) invasion by parasites (*CR1, GRK5*) or RBC defects. Among those, sickle cell trait (haemoglobin S, *HbS*), haemoglobin C (*HbC*) at homozygote state, alpha+ thalassemia and glucose-6-phosphate dehydrogenase (*G6PD*) deficiency have been associated with a protection against parasite invasion and clinical malaria attacks *(7)*.

Other association studies have focused on chromosomal regions linked with parasite infection levels and mild malaria susceptibility, in particular 5q31-33 and 6p21-23 (8–15). In this last region, *TNF* has been the most studied candidate; *NCR3*, encoding a cell membrane receptor of natural killer, has been also repeatedly associated with mild malaria (10,16).Since 2010, several genome-wide association studies (GWAS) and meta-analysis have been published on severe malaria (17–21). These studies confirmed the involvement of previously known susceptibility genes (*HBB* and *ABO*) and revealed new genes (*ATP2B4*, the cluster of genes *GYP/FREM3*, among which *GYPA* and *GYPB* appear to play a central role (21)).

Here we present the first GWAS performed on mild malaria susceptibility. It was conducted on two cohorts of infants in southern Benin, used as discovery and replication cohorts (525 and 250 individuals respectively) and genotyped with the high density Illumina^®^ HumanOmni5 beadchip. This study intended to identify genetic factors that play an early role in disease development (Figure 1), in cohorts of infants followed from birth to 18-24 months of age, thus targeting factors involved in innate immunity or innate resistance mechanisms.

**Figure 1.**
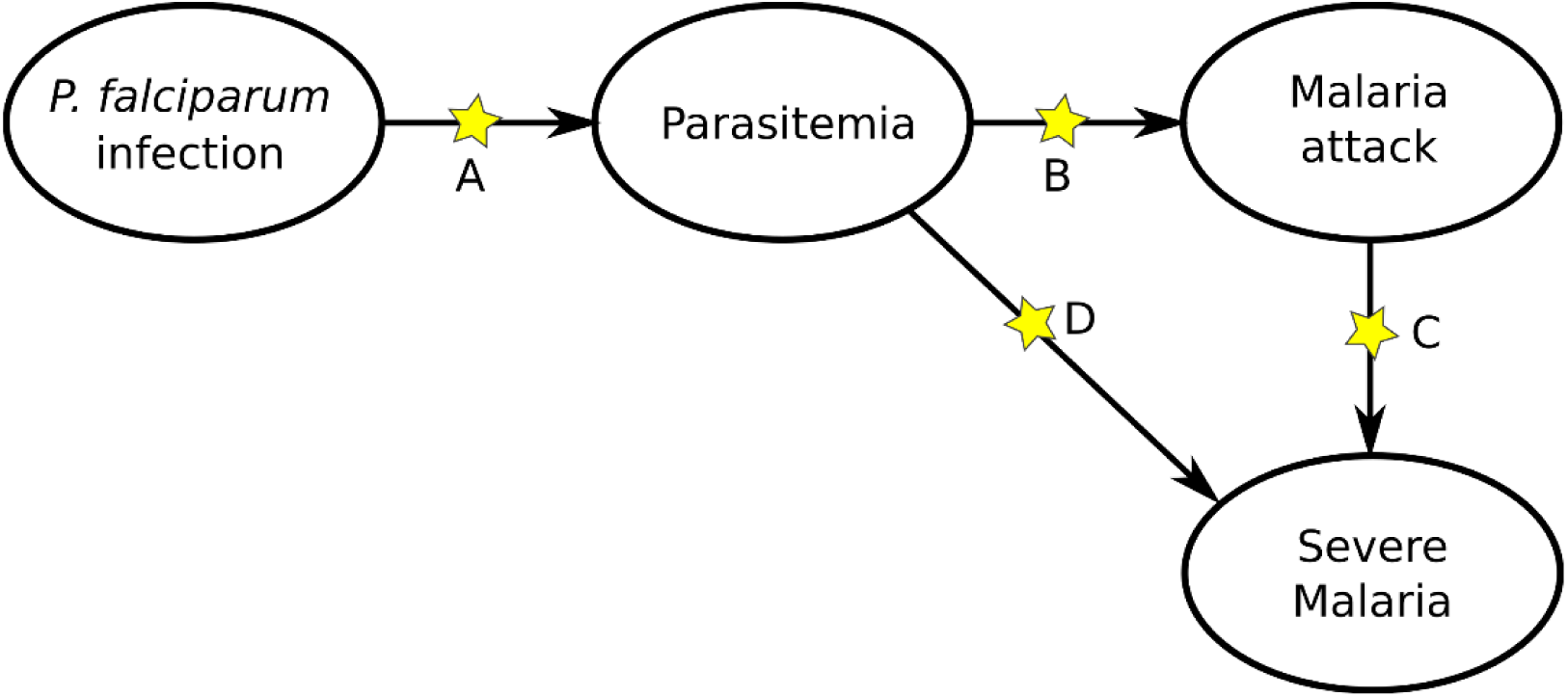
The path from *P. falciparum* infection to severe malaria, through parasitemia and malaria attacks (parasitemia with fever). The process may stop at each transition A, B, C and D, depending on several factors including the genetics of the host. Studies on severe malaria mainly target factors involved in transitions C and D, while studies on mild malaria focus on A and B. Our study focuses on transitions A and B in a population of infants.

In our study, infants were closely followed with symptomatic and asymptomatic malaria infections recorded. Furthermore geographical, climatic, behavioral and entomological data were collected in concomitant environmental risk surveys throughout the follow-up, to estimate a spatial-and time-dependent risk of exposure. Association studies were performed with the recurrence of mild malaria attacks (RMM) and the recurrence of malaria infections (RMI) including mild malaria attacks and asymptomatic infections, taking into account a time-dependent risk of exposure. We find convincing evidence supporting the involvement of *PTPRT, MYLK4, VENTX* and *UROC1* as well as strong association signals in *PTPRM, ACER3 and CSMD1* all of which are relevant putative candidate genes in malaria physiopathology.

## Results

### Phenotypic and genotypic data

Infants from both discovery (Tori-Bossito) and replication (Allada) cohorts come from southern Benin, from two sites located 20 km from each other, 40-50 km north-west of Cotonou, the economic capital. Ethnicity composition, based on self-reported ethnicity of the mother, significantly differed between the two study sites. The main ethnic groups were Tori (74%) and Fon (11%) in the first cohort whereas they were Aïzo (68%) and Fon (22%) in the second one. Each of the other ethnic groups accounted for less than 10% of individuals in each cohort.

Principal Component Analysis (PCA) performed on the two samples did not reveal any ethnic outliers and showed a low degree of stratification (Figure S1). PC1 separated the two cohorts and PC2 likely reflected some heterogeneity in the Tori-Bossito cohort. We observed that infants did not cluster by reported ethnicity, but rather by cohort and health center, indicating that the self-reported ethnicity of the mother is not a major factor of genetic heterogeneity in our samples. The PCA including 1000 Genomes Project (KGP) populations (as described in methods) confirmed that our two cohorts are homogeneous (Figure S2). As expected, our two populations overlapped and clustered with Nigerian populations (Yoruba and Esan).

Main characteristics of infants were compared between samples included in or excluded from GWAS (Tables S1-S2). For the discovery cohort, no significant differences were observed except for ethnic group composition which appeared to be enriched in Tori at the expense of less prevalent ethnic groups (p<0.04). For the replication cohort, only infants followed between 12 and 24 months of age were included in the GWAS. Thus a higher proportion of infants were excluded from analyses (infants lost to follow-up during the first year). We observed in our sample a lower proportion of infants with low birth weight and born to mothers who reported having never attended school (p<0.01 and p=0.06 respectively).

For the 525 infants included in the discovery sample, mean (SD) length of follow-up was 16.9 months (2.83). A total of 342 infants (65.1%) experienced at least one mild malaria attack (from one to 10 episodes) and 359 infants (68.4%) were observed with at least one malaria infection (range from one to 16 malaria infections by infant). For the replication cohort of 250 infants, mean (SD) length of follow-up was 11.9 months (1.72). A proportion of 83.2% and 86.8% children experienced at least one malaria attack (range 1 to 9), or at least one malaria infection (range 1 to 14) during the follow-up, respectively.

### Adjusting on main epidemiological and environmental determinants

Genome-wide association analyses were performed in two steps because computational burden inherent to the random effect Cox model (23) makes it inappropriate for the large number of tests in GWAS.

In the first step, the recurrence rate of malaria episodes was modeled using a random effect Cox model for recurrent events, taking into account epidemiological and environmental factors that might influence malaria infections in infants. A few covariates among all those considered were retained in the final model (Tables S3-S4). In each cohort, the same covariates were found associated for the recurrence of mild malaria attacks (RMM) and the recurrence of malaria infections (RMI). The levels of exposure to vector bites, health centers and transmission seasons were associated with the risk of infection in both cohorts. In addition, for the Allada cohort, significant effects were observed for marital status and for maternal education level. No effects of placental infection (detected by a thick and thin placental smears), low birth weight (<2500 g) or HbS allele carriage were detected.

### Genome-wide association analyses

After imputation using the KGP reference panel (phase 3), 15,566,900 high quality (R^2^ > 0.8) variants with a minimum allele frequency (MAF) ≥ 0.01 were available for analysis. Association was tested at a genome-wide level using a linear mixed model, with confounder-adjusted phenotypes constructed from the model built at the previous step. These adjusted phenotypes correspond to individual effects which are not explained by the epidemiological covariates. The genomic inflation factor was consistent with a reliable adjustment for cryptic relatedness in our samples, for both analyses (Figure 2).

**Figure 2.**
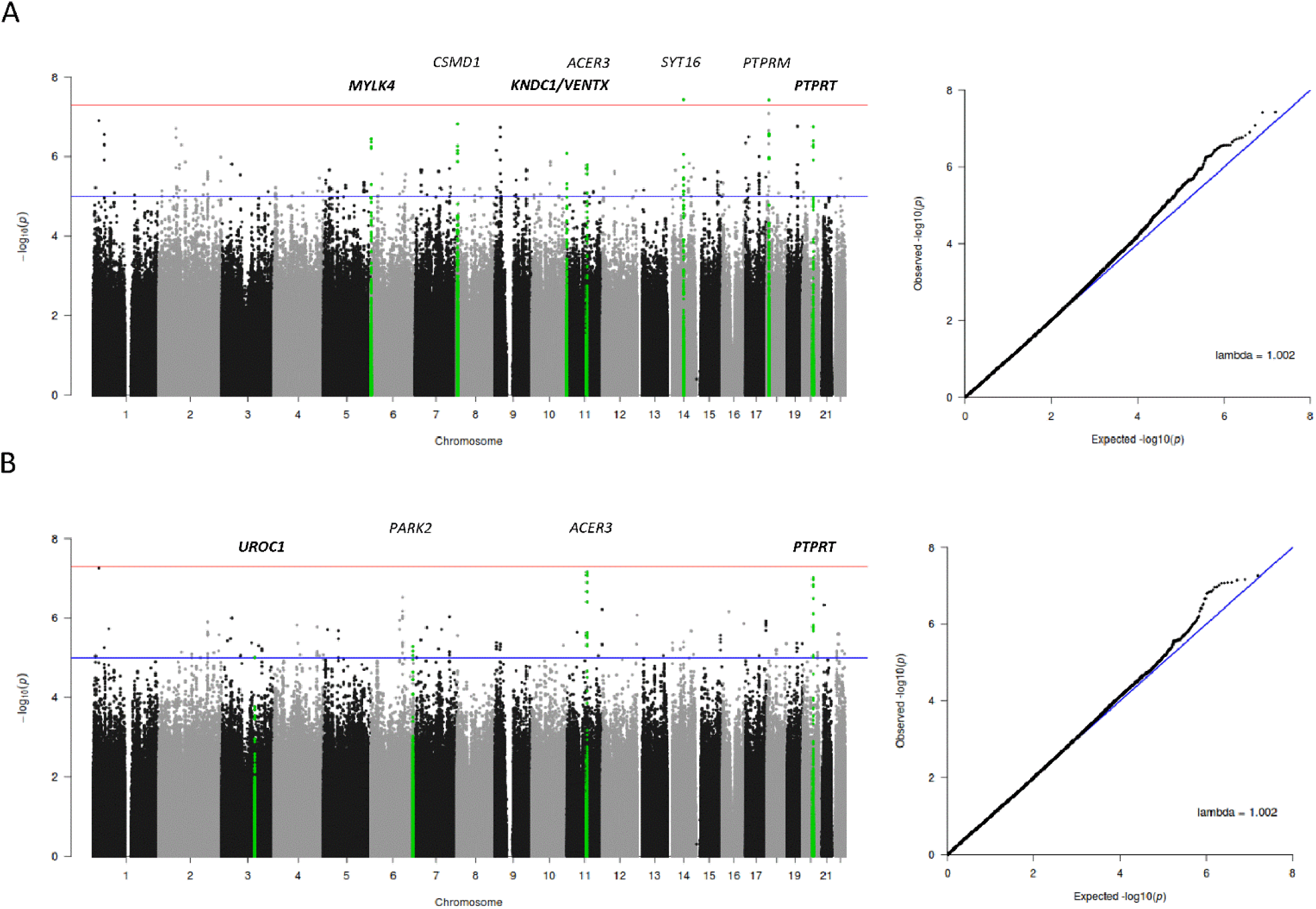
Manhattan plots and QQ plots for the discovery association study. A) GWAS results for the recurrence of mild malaria attacks, B) GWAS results for the recurrence of malaria infections. The red line in the Manhattan plots indicates the threshold of significance (5 × 10^−8^), the blue line the threshold of suggestive association used for replication (1 × 10^−5^). The blue line in the QQ plots shows the 1:1 regression line (the expected distribution of p-value under the null hypothesis). Lambda is the genomic inflation factor calculated as the ratio of the median of the empirically observed distribution of the test statistic to the expected median.

The analysis revealed four association signals with a *p*-value very close to the genome-wide association threshold of 5×10^−8^: two signals for RMM (Figure 2A), located in *SYT16* (14q23.2 region, lead SNP rs375961263, p=3.7×10^−8^) and in *PTPRM*, a gene encoding a receptor-type protein tyrosine phosphatase (18p11.23 region, lead SNP rs113776891, p=3.77×10^−8^); and two signals for RMI (Figure 2B): one located in *ACER3*, a gene encoding an alkaline ceramidase (11q13.5 region, lead SNP rs77147099, p=6.85×10^−8^) and another one located in *PTPRT*, a gene encoding a second receptor-type protein tyrosine phosphatase (20q12 region, lead SNP rs111968843, p=9.70 ×10^−8^).

For these signals, data were re-analyzed in a single step with a mixed Cox model for recurrent events accounting for cryptic relatedness and relevant covariates, without resorting to intermediate confounder-adjusted phenotypes. Minor differences were observed in p-values and none of them reached genome-wide significance.

A total of 356 and 214 variants were associated at *p*<1×10^−5^ with RMM and RMI respectively (Tables S5-S6), and were tested for replication in the second cohort. SNPs that showed evidence of replication (*p*<0.05) are presented in Tables 1 and 2. Highest statistical support was observed for SNPs located in the 6p25.2 locus with RMM (3 SNPs, min-*p* = 0.0056). The most strongly associated SNP in the replication cohort, rs72840075, is located in an intron of *MYLK4* which encodes a myosin light chain kinase (Figure 3). A meta-analysis of these SNPs showed a borderline significant association for SNP rs142480106 (*p*=5.29 × 10^−8^) (Table 1).

**Table 1.** Association signals which replicated at *p* < 0.05 for the recurrence of mild malaria attacks.

| Locus | SNP | Risk allele | MAF | $p$<br>initial<br>GWA<sup>a</sup> | $p$<br>Cox model<br>Tori-bossito<sup>b</sup> | HR (CI 95%)<br>Tori-Bossito | $p$<br>Cox model<br>Allada<sup>b</sup> | HR (CI 95%)<br>Allada | Nearest Genes | $p$<br>Meta-analysis |
| --- | --- | --- | --- | --- | --- | --- | --- | --- | --- | --- |
| 6p25.2 | rs76088706 | T | 0.025 | $5.56 \times 10^{-7}$ | $2.68 \times 10^{-6}$ | 2.22 (1.59-3.11) | 0.036 | 1.75 (1.03-2.96) | C6orf195(Intergenic) | $3.65 \times 10^{-7}$ |
| | rs142480106 | T | 0.019 | $4.35 \times 10^{-7}$ | $1.14 \times 10^{-6}$ | 2.52 (1.73-3.66) | 0.013 | 2.08 (1.17-3.70) | MYLK4 (UTR3) | $5.29 \times 10^{-8}$ |
| | rs72840075 | A | 0.030 | $5.04 \times 10^{-6}$ | $1.05 \times 10^{-5}$ | 2.03 (1.48-2.79) | 0.0056 | 1.88 (1.20-2.95) | MYLK4 (intronic) | $2.01 \times 10^{-7}$ |
| 10q26.3 | rs182416945 | A | 0.12 | $8.31 \times 10^{-7}$ | $5.56 \times 10^{-6}$ | 1.55 (1.28-1.87) | 0.02 | 1.28 (1.04-1.58) | KNDC1 (intronic) | $7.94 \times 10^{-7}$ |
| 20q12 | rs6124419 | A | 0.12 | $1.21 \times 10^{-6}$ | $2.51 \times 10^{-6}$ | 1.53 (1.28-1.83) | 0.04 | 1.26 (1.01-1.58) | PTPRT (intronic) | $6.82 \times 10^{-7}$ |

HR, hazard ratio; MAF, minor allele frequency
<sup>a</sup>p-value in GWA analysis performed in two steps using a confounder-adjusted phenotype.
<sup>b</sup>p-value of the association calculated with a random effect Cox model accounting for cryptic relatedness and relevant covariates (health center, risk of exposure, transmission season for both cohorts, marital status and education of women in addition for Allada cohort)

**Table 2.**
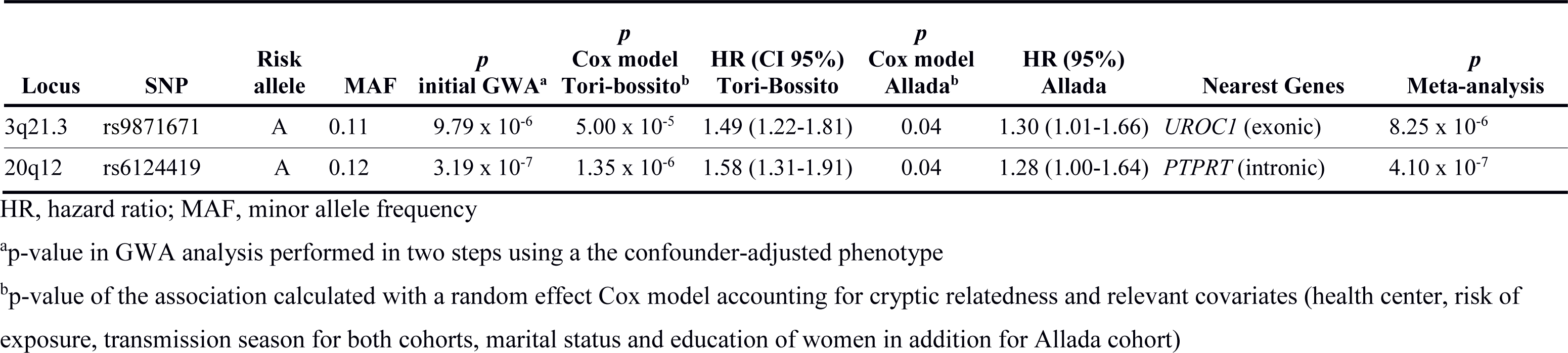
Association signals which replicated at p < 0.05 for the recurrence of malaria infections.

**Figure 3.**
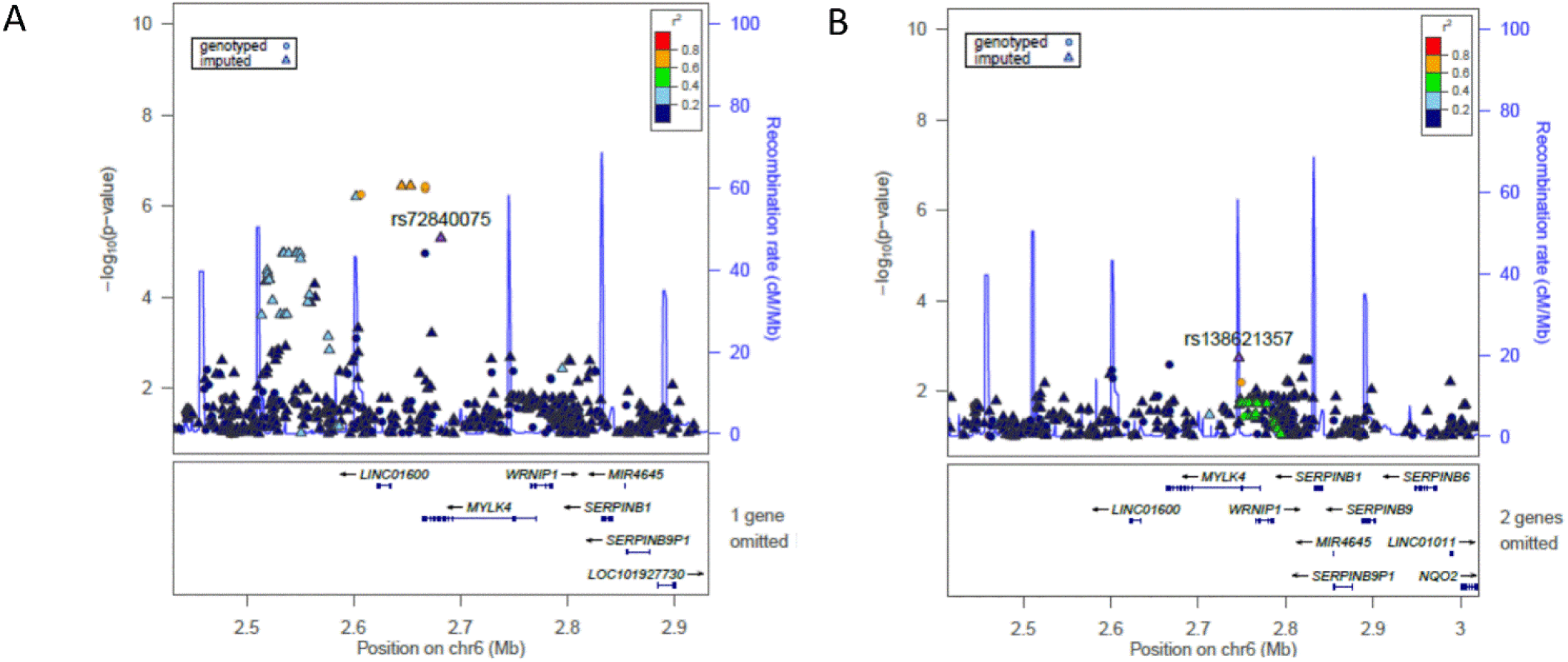
Annotated regional association plot for the 6p25.2 locus: A) Association results with the recurrence of mild malaria attacks, pairwise LD are shown with rs72840075, the best associated SNP in replication analysis B) Association results in conditional analysis on rs72840075. Plots show LD calculated on Nigerian Population of KGP dataset (ESN and YRI populations) in the 250 kb region.

The same SNP, rs6124419, located in intron 18 of *PTPRT*, replicates for both phenotypes. The regional linkage disequilibrium (LD) plot showed a narrow signal of around 20 kb width in both cases which encompasses exons 20 to 22 (Figure 4A). An estimation of the incidence rate of mild malaria attacks in three risk groups based on exposition levels showed that the effect of rs6124419 is consistent regardless of the exposition level and the cohort (Figure 4C). Interestingly *PTPRT* is a paralogue of *PTPRM*, for which a highly suggestive association peak was observed with mild malaria attacks in the 18p11.23 chromosomal region (Figure 4A).

**Figure 4.**
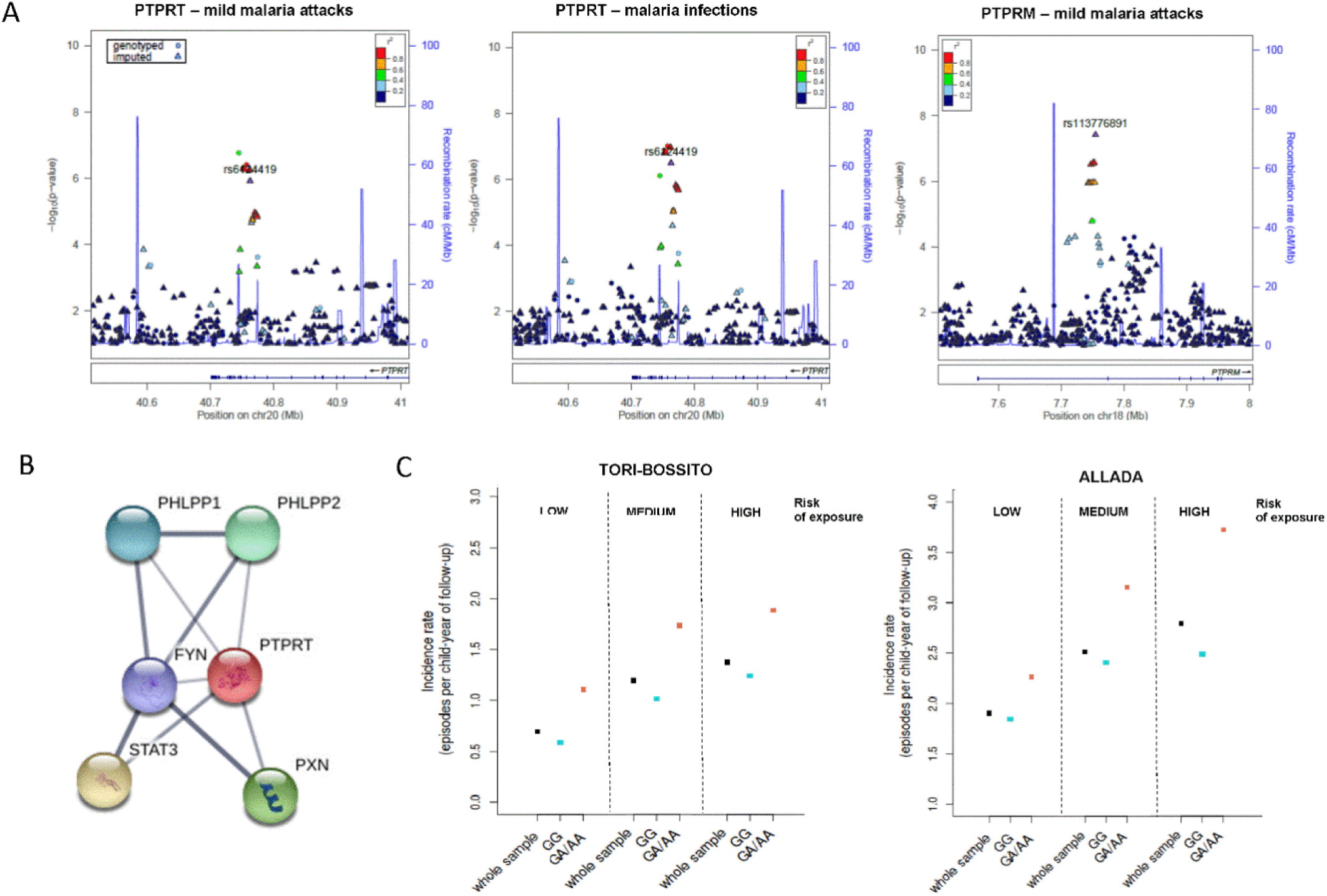
A) Annotated regional association plots for *PTPRT* and *PTPRM* loci. B) Interaction network for PTPRT C) Crude incidence rate of mild malaria attacks for rs6124419 genotypes calculated for three classes of environmental exposure. Plots show LD calculated on Nigerian Population of KGP dataset (ESN and YRI populations). Replicating SNP are highlighted for PTPRT. The three classes were defined from the mean environmental exposure risk across all the follow-up calculated for each infant. Low, medium and high correspond to the three tertiles of the distribution.

Other associations which replicated in the Allada cohort include rs182416945 in *KNDC1* and rs9871671 in *UROC1* (Table 2). This last one, a missense SNP, is predicted to be probably damaging with a score of 0.99 in PolyPhen-2*(24)* and has a Combined Annotation Dependent Depletion (CADD)*(25,26)* score of 26.5, indicating that this variant is among the 1% most deleteriousness substitutions of the human genome.

For each signal aforementioned, a regional LD plot for the nearby region (± 200 kb) was realized and a conditional analysis was run (Figure 4A and Figures S3 and S4). For none of them, a significant residual signal was found.

### Functional annotation and gene mapping of GWAS results

To identify the genes and variants most likely associated with these four replicating association signals, and to select potential genes of interest pointed out by non-replicating signals, we used the FUMA web platform *(27)*. FUMA identifies independent associated genomic regions and prioritizes genes based on functional consequences of SNPs in each of them.

Thirty independent associated regions were identified for RMM and 15 for RMI, based on the presence of at least 3 SNPs below the *p*-value threshold of 10^−5^ in a region (LD threshold to aggregate SNPs in one signal, r^2^ = 0.1). Candidate SNPs (among SNPs present either in our data or in the AFR population of the KGP reference panel) were then defined based on LD (r^2^>0.6) with the lead SNP in each of these regions, or with one of the replicating SNPs.

These candidate SNPs were mapped to genes, by positional mapping (deleterious coding SNPs - CADD score > 12.37 – or regulatory elements likely to affect binding in non-coding regions - Regulome database (RDB, (28)) score ≤ 2), and by eQTL mapping based on expression data from relevant tissue types for malaria (whole blood, cell-EBV transformed lymphocytes, liver, skin, cells transformed fibroblasts and spleen). These mappings identified 20 genes for RMM and 8 for RMI (Tables S7-S8). Results for the strongest signals of association with both phenotypes are summarized in Figure 5. We additionally performed chromatin interaction mapping with high chromosome contact map (Hi-C) data (29) with FUMA, allowing to identifying genes located in regions with significant chromatin interaction with the candidate SNPs (Figure 5).

**Figure 5.**
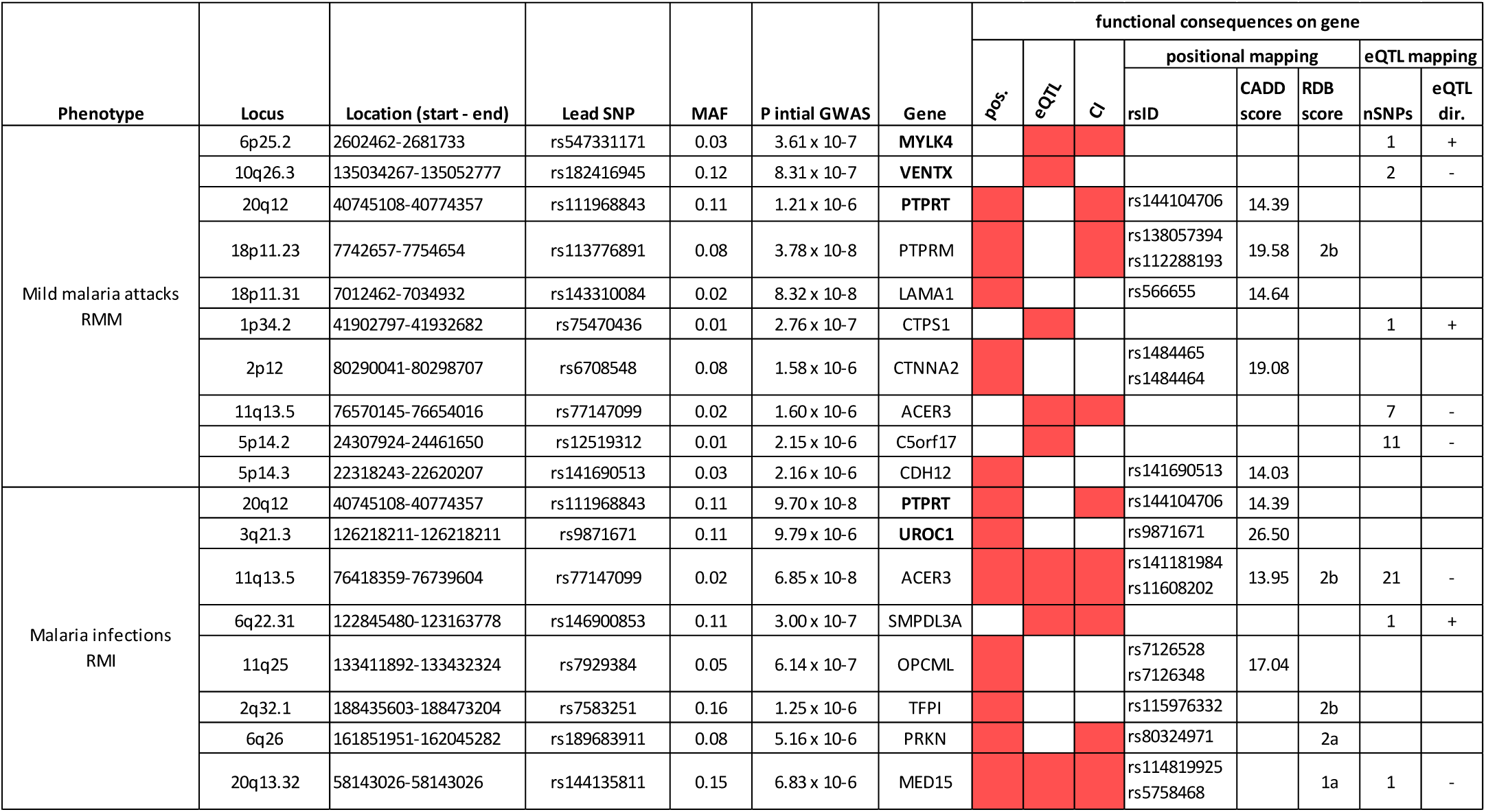
Genes identified by FUMA based on functional consequences of SNPs, in replicating association signals and in the strongest non-replicating ones. Genomic regions reported are those with at least one gene identified based on positional mapping or eQTL mapping. Genes highlighted in bold correspond to replicating signals. For each genomic region the lead SNP was identified as the replicating SNP or the SNP with the lowest *p*-value. The *p*-value of the initial GWA was calculated with a linear mixed model on the adjusted phenotypes. Filled red boxes indicate whether the gene was identified by positional mapping (pos.), eQTL mapping (eQTL) or chromatin interaction mapping (CI). For positional mapping, deleterious SNP names are given (rsID) with the maximum CADD score and RegulomDB score observed for these SNPs. For eQTL mapping, the number of significant eQTL associations (nSNPs) are reported, together with the direction of effect allele to gene expression, for the risk increasing allele of GWAS (eQTL direction).

For each of the four replicating loci FUMA identified a single gene, suggesting that these genes are very likely to be the “causal gene” in these regions. Three of these highlighted genes correspond to the gene in which the replicating SNP is located: *MYLK4* in 6p25.2 region, *PTPRT* in 20q12 region, *UROC1* in 3q21.3 region. The last highlighted gene, VENTX in 10q26.3 region, is located 17kb from the lead SNP. In *MYLK4* locus, the replicating SNP, rs72840075, is an eQTL in whole blood (eQTLGen and BIOS eQTL (30) database, FDR<0.05). Allele at risk in our data is associated with a higher expression of this gene. One deleterious SNP is observed in *PTPRT* (r^2^=0.78 with the replicating SNP, CADD = 14.39), and in *UROC1* (the aforementioned deleterious SNP). In *VENTX* locus, two SNPs, rs182416945 (the replicating one) and rs138609386, in complete DL (r^2^ = 1) with the first one, were identified as a significant eQTL in whole blood (eQTLGen database). The allele at risk in our data is associated with a lower expression of *VENTX*, a gene coding for the VENT Homeobox.

FUMA highlighted also *PTPRM* as the most probable gene associated with the prominent signal for RMM in 18p11.23 region. Two candidate SNPs, rs138057394 and rs112288193 (r^2^=0.74 with the lead SNP, for both SNPs) are annotated as deleterious coding SNPs (CADD score of 19.58 and 14.64 respectively).

Furthermore, among the other genes pointed out by FUMA (Figure 5) in non-replicating signals, *ACER3* in 11q35.5 region is identified by the three mapping methods performed. Numerous significant eQTL associations were observed in the cells transformed fibroblast tissue (GTEx v7 database). For these eQTL, alleles at risk in our data were associated with a lower expression of the gene. This association signal is the strongest signal for RMI, with a *p*-value approaching the genome-wide significance threshold, (*p* = 6.85 × 10^−8^, figure 2) and it was also found associated with RMM (*p* = 1.60 × 10^−6^).

### Candidate associations

We also tested the statistical significance of genes and variants previously reported in literature as associated with malaria (Table 3). Only SNP rs40401 in *IL3* was found marginally associated with both phenotypes (p = 0.01 and 0.02 respectively). Moreover, two suggestive peaks are observed in the candidate regions: one in the *IL3* locus with both phenotypes (10 SNPs encompassing *IL3* and exon 1 to 3 of *ACSL6*, min-p = 8.89×10^−5^ with RMM and min-p=1.66×10^−4^ with RMI, rs7714191) and one in the *IL10* locus with RMM, 26 kb upstream of the gene (4 SNPs, min-p = 2.04×10^−4^, rs116126622).

**Table 3.** Candidate SNP associations

| Gene | SNP | Variant | Location<br>(GRCh37) | MAF | Change | Malaria attacks |  | Malaria infections |  |
| --- | --- | --- | --- | --- | --- | --- | --- | --- | --- |
| | | | | | | $\beta$(CI 95%) | p | $\beta$(CI 95%) | p |
| *HBB* | rs334 | HbS | 11:5248232 | 0.10 | A > T | -0.040 | 0.24 | 0.05 | 0.34 |
|  | rs33930165 | HbC | 11:5248233 | 0.04 | G > A | -0.011 | 0.83 | -0.01 | 0.95 |
| *TNF* | rs1799964 | 3'UTR, *TNF* -857 | 6:31574531 | 0.19 | T > C | 0.002 | 0.92 | 0.02 | 0.59 |
|  | rs1800629 | 3'UTR, *TNF* -308 | 6:31543031 | 0.11 | G > A | 0.018 | 0.11 | -0.011 | 0.80 |
| *IL10* | rs1800871 | 3'UTR, *IL10* -819 | 1:206946634 | 0.38 | C > T | 0.023 | 0.23 | 0.02 | 0.51 |
| *IL3* | rs40401 | Exon 1, missense | 5:131396478 | 0.71 | C > T | 0.050 | 0.01 | -0.08 | 0.02 |
| *ABO* | rs8176746 | Exon 10, missense | 9:136131322 | 0.15 | C > A | -0.022 | 0.42 | -0.062 | 0.13 |
|  | rs8176719 | Exon 9, frameshift | 9:136132908 | 0.27 | - > G | -0.027 | 0.18 | -0.038 | 0.22 |
| *ATP2B4* | rs1541255 | Exon 1 | 1:203652141 | 0.41 | A > G | -0.006 | 0.75 | -0.035 | 0.24 |
|  | rs10900585 | Intron 2 | 1:203654024 | 0.46 | G > T | 0.000 | 0.96 | 0.014 | 0.65 |

MAF, minor allele frequency;
Results of association tests performed in the discovery cohort using the confounder-adjusted phenotype. $\beta$ is the estimate of the SNP effect under an additive model. A negative value indicates a protective effect of the alternative allele and conversely.

## Discussion

Genetic factors of resistance to malaria may be involved at different stages during the course of the disease: inoculation of parasite, development of asymptomatic parasitemia, appearance of clinical symptoms including fever, rapid progress to complications and manifestation of severe malaria (4). However research efforts to identify host genetic factors have so far mainly focused on severe malaria, the life threatening form of the disease. While some genetic factors may be in common between severe and mild forms, studies on severe malaria are not designed to discover genes affecting specifically the risk of malaria infection and the development of mild malaria. For this reason, studies based on non-severe malaria forms may help to complete our understanding of the disease.

Our study represents the first screening of genetic variations associated with non-severe forms of malaria at a genome-wide level. It was conducted on two birth cohorts closely followed with a protocol of surveillance which allowed to ascertain malaria infections as a whole. Infants were visited at home weekly or bi-monthly during all the follow-up for the systematic detection of fever and symptomatic malaria. Moreover, a TBS was performed monthly to detect asymptomatic infections (23,24). Another strength of our study is the assessment of environmental risk of exposure at an individual level all along the follow-up. Although there is consistent evidence for local variation in exposure to *P. falciparum*, which may partly explain the heterogeneity in malaria infection incidence observed in children (31,32), this factor has been seldom considered in genetic epidemiology studies. Here, environmental risk of exposure was taken into account in an unprecedented manner. Entomological, climatic, environmental (information on house characteristics and on its immediate surrounding) as well as geographical data (soil type, watercourse nearby, vegetation index, rainfall, etc) were collected all along the follow-up. Altogether these data allowed modeling, for each child included in the follow-up, an individual risk by means of a time-dependent variable (33). The benefit of using this estimated risk has been demonstrated in two studies on the impact of placental malaria on infant susceptibility (34,35). This risk turned out to have a highly significant effect on the recurrence of events in the present study (p<1×10^−8^ and p<1×10^−6^ for discovery and replication cohort respectively).

We find strong support for the involvement of protein tyrosine phosphatase (PTP) receptors. Highly suggestive association peaks were observed for *PTPRM* with RMM and association peaks observed for *PTPRT* replicated for both phenotypes. Interestingly, both genes are paralogues and in each signal a deleterious coding SNP which could be the causal SNP is identified. Although the replication of *PTPRT* was weak in the second cohort (p=0.01), this can be explained by the fact that our replication cohort is relatively small in size, that infants in this cohort were not actively followed during the first year of life, which limited our analysis to the 12-24 months period. However, assessment of incidence rate of mild malaria attacks in function of groups of different level of exposure, showed very consistent results among exposure groups and cohorts which is an additional argument for a true association. PTPRT and PTPRM are proteins belonging to the PTP superfamily, which regulates diverse signaling pathways by catalyzing the removal of a phosphate group from specific signaling proteins. PTPRT and PTPRM encodes two PTP receptors of the same sub-family type IIb *(36)*. These transmembrane proteins have an extracellular domain involved in cell-cell aggregation and a phosphatase domain which allows intra-cellular signaling. Interestingly PTPRT protein directly interacts with STAT3 as shown by the STRING interaction Network (Figure 4 B) *(37)* and is the only interactor for which a high confidence (>0.70) is reported in the STRING database. The role of PTPRT in regulating STAT3 pathways has been demonstrated, with STAT3 identified as a direct substrate of PTPRT (38,39). STAT3 is a signal transducer and activator of transcription which behaves similarly to NF-κB, the role of which in malaria infection is well-established. Moreover, STAT3 has been reported as associated with cerebral malaria severity in experimental studies (40,41). Liu et al. have demonstrated using murine models that *STAT3* is activated by *P. berghei* infection. The STAT3 pathway is thus very likely to be involved in the first stages of malaria infection and represents a potential drug target.

Other genes replicated with p *<* 0.05 in the validation cohort. *MYLK4* belongs to the family of myosin ligth chain kinases (MYLKs) that catalyse myosin interaction with actin filaments. Since members of MYLKs family play an essential role in the organization of the actin/myosin cytoskeleton, and in cell motility (42), *MYLK4* may play a role in RBC membrane structure. At 10q26.3 locus, replication is observed for a SNP located in *KNDC1*, but two eQTL associations identified *VENTX* as the most probable effector for this association signal. *VENTX* is a homeobox transcriptional factor that controls proliferation and differentiation of hematopoietic and immune cells (43–45). Wu et al. works (44,45) have shown that *VENTX* is a key regulator of macrophage terminal differentiation and of dendritic cells differentiation and maturation. *VENTX* is up-regulated during monocyte-to-macrophage differentiation and overexpression of *VENTX* accelerates the differentiation towards dendritic cells. Conversely, *VENTX* deficiency impairs dendritic cells differentiation and affects various aspects of macrophage differentiation and function, including a downregulation of multiple membrane receptors critical for innate and adaptive immunity. Both macrophages and dendritic cells are innate immune cells derived from monocytes that play an essential role in the response to malaria infections (Chua et al., 2013). Allele at risk of mild malaria attacks in our data was associated with a lower expression of *VENTX*, which would negatively impact the macrophages and/or dendritic cells response to malaria infection in the most susceptible children. Finally, a high deleterious missense SNP in *UROC1 (*Urocanate Hydratase 1), a gene expressed in the liver (GTEX RNA-seq), is also associated. Urocanate Hydratase or urocanase is an enzyme that catabolizes urocanic acid to 4-imidazolone-5-propionic acid in liver. Urocanic acid is found in the skin and the sweat of humans, and has been shown to protect the skin from ultra violet radiation. It has also been demonstrated to be a major chemo-attractant for a skin-penetrating parasitic nematode (46). Thus, urocanic acid and urocanase could be hypothesized to play a role in the attractiveness of humans to mosquitoes.

Several strong association signals in the discovery cohort were not replicated in the validation cohort. Given the limits of the replication cohort mentioned above, we think that the most promising ones are still worth consideration and functional annotation and gene mapping at these loci were performed with FUMA. *ACER3* identified at 11q13.5 locus appears of particular relevance. It is associated both with a protection against asymptomatic and symptomatic malaria infections. *ACER3* is an alkaline ceramidase, involved in the degradation of ceramides into sphingosine-1-phosphatase (S1P). Ceramides have an anti-plasmodium action (47,48), which is inhibited by S1P; and it has been hypothesized that antimalarial drug artemisinin and mefloquine have an antiparasitic action through the activation of sphingomyelinase producing ceramide (49). In our data allele at risk is associated with a decreased *ACER3* expression, thus indicating a lower ceramidase activity in the children with higher recurrence of malaria infections. This is not in accordance with the anti-plasmodium action of ceramides as mentioned above as a lower *ACER3* expression would be rather associated with a higher availability of ceramides. However *ACER3* has been also involved in the activation of innate immune in mice and this could also to explain the role of ACER3 observed in malaria protection *(50)*.

A second gene of interest highlighted by FUMA in malaria infections analyses is *PRKN* (or PARK2) coding for the parkin RBR E3 Ubiquitin Protein Ligase. Mutations in this gene are known to cause Parkinson disease and this gene has also been involved in innate immune response with a role in ubiquitin-mediated autophagy of intracellular pathogens, specifically *M.tuberculosis (51)*. Finally, a strong association signal in *CSMD1* gene on chromosome 8 was evidenced in mild malaria attacks analysis. No functional evidence was found by FUMA for this association signal; however, an association signal in *CSMD1* was recently found in a GWAS of severe malaria in Tanzania (20); it is a transmembrane protein, assumed to be a tumor suppressor gene (52), and is also associated with phenotypes as diverse as blood pressure (53) and schizophrenia (54) but there is as of yet no clear scenario to explain its potential role in malaria.

We also examined association results for the most associated candidate variants with malaria-related phenotypes in the literature (Table 3). There was no evidence of replication for any variant, except rs40401 located in *IL3* (p = 0.01 and p = 0.02 for malaria attacks and malaria infections, respectively). The fact that our study target population differs substantially from those used in most published studies on mild malaria may partly explain the limited evidence of replication observed. *HBB* was not significantly associated with non-severe malaria in our sample. There are accumulating evidence for an age-related protective effect of HbS (55–57) with an acquired mechanism of protection from both asymptomatic and symptomatic *P. falciparum* infections in children. Our results are in line with two of these studies (55,56) which did not detect a protective effect before the age of two.

Our study has identified several genes whose biological function is relevant to malaria physiopathology and which could play a role in the control of malaria infection. Naturally, future studies are required to further validate these findings. The difficulty of setting up such longitudinal field studies have indeed limited the number of individuals that have been included in malaria cohorts. However our results show that GWAS on non-severe malaria can successfully identify new candidate genes and inform physiological mechanisms underlying natural protection against malaria. Improving our understanding of the disease course is crucial for the development of effective control measures.

## Materials and Methods

### Study population – discovery cohort

The discovery cohort was composed of 525 infants followed-up from birth until 18 months (58). This study took place from June 2007 to January 2010 in 9 villages of Tori-bossito district located 40 km North-East of Cotonou. Southern Benin is characterised by a subtropical climate, with 2 rainy seasons (a long rainy season from April to July and a short one in August and September). Clinical incidence of malaria mainly due to *P. falciparum* (97%) was estimated to be 1.5 (95%CI 1.2-1.9) malaria episodes per child (0-5 years) per year in this area (59) and entomological inoculation rate on average 15.5 infective bites per human per year in studied villages (33). The design and the flow-chart of the study have been published elsewhere (58). Briefly, 656 infants were included at birth and followed with a close parasitological and clinical survey until the age of 18 months. During the entire follow-up, infants were weekly visited at home by a nurse of the program and temperature was systematically controlled. In case of axillary temperature higher or equal to 37°5 (or a history of fever in the preceding 24 hours), child was referred to health center for medical screening where both a TBS and a rapid diagnostic test (RDT) were performed. Once a month a systematic TBS was performed to detect asymptomatic parasitemia. At any time in case of suspicious fever or clinical signs, related or not to malaria, mothers were invited to bring their infants to the health center where the same protocol (temperature, TBS and RDT) was applied. Symptomatic malaria infection was treated with artemether-lumefantrine combination therapy as recommended by the National Malaria Control Program. A total of 10589 TBS were performed along the follow-up which were read by two independent technicians.

To assess the risk of exposure to *Anopheles* bites, environmental (information on house characteristics and on its immediate surrounding) and geographical data (satellite images, soil type, watercourse nearby, vegetation index, rainfall, etc) were recorded. Throughout the study, every six weeks, human landing catches were performed in several points of the villages to evaluate spatial and temporal variations of *Anopheles* density. Altogether these data allowed modeling, for each child included in the follow-up, an individual risk of exposure by means of a space-and time-dependent variable (33).

### Study population – replication cohort

The replication cohort was composed of 250 infants who were part of a mother-child study conducted from 2009 to 2013 in the district of Allada, a southern semi-rural area located 55 kilometres north of Cotonou and 20 km from the first study site (60). Infants were born to mothers who participated in a multi-country clinical trial for the prevention of malaria in pregnancy (MiPPAD trial, “Malaria in Pregnancy Preventive Alternative Drugs, NCT00811421,(61)). The first 400 newborns from MiPPAD participants (62) were enrolled in the present study. In brief, 400 offsprings were included at birth between January 2010 and June 2012 and among them 306 integrated the second year follow-up. During the first year, detection of malaria cases was passive Children were seen three times at 6, 9 and 12 months of age for scheduled visit. At these occasions a TBS was performed for malaria screening. Between 12 and 24 months the same protocol of follow-up as in the first study was set up. Infants were visited at home every two weeks by a nurse and temperature was taken. As before in case of axillary temperature greater than or equal to 37.5°C, child was referred to health center where a RDT and a TBS were performed. Each month a systematic TBS was made to detect asymptomatic malaria. During the whole 24 months follow-up period, in case of suspicious fever or any health problems, mothers were invited to visit health centers where the same protocol for malaria screening was applied (temperature, TBS and RDT). Clinical malaria infections were treated as recommended by the National Malaria Control Program. Environmental and geographical data were collected as well, to estimate risk of exposure for each infant. The methodology detailed in Cottrell et al., was used again except that this time *Anopheles* density was measured in the children’s room, using a CDC light trap.

In both cohorts, children included in analysis had a minimum of three months follow-up. For the replication study, we choose to consider only the second year follow-up because i) the differences of malaria follow-up protocols in the first year lead to the detection of a lower number of malaria infections, ii) environmental and entomological data needed to estimate the risk of exposure were missing for children who did not enter the second year follow-up. A description of main characteristics of infants included or not in the GWAS are presented in supplementary table S1 and S2.

### Ethics Approval

For the two cohort studies, both oral and written communications were provided to parent’s children interested in participating. An informed consent, written in French and in Fon, was presented to, and signed by parents who agreed to participate. The protocols of these studies were approved by both the Beninese Ethical Committee of the Faculté des Sciences de la Santé (FSS) and the IRD Consultative Committee on Professional Conduct and Ethics (CCDE).

### Genotyping and data quality control

Genotyping was conducted at the Centre National de Recherche en Génomique Humaine (CNRGH, CEA, Evry, France). Before genotyping, a quality control was systematically performed on each DNA sample. All samples were quantified by fluorescence, in duplicate, using the Quant-It kits (Thermo Fischer Scientific). The lowest values systematically underwent a second measurement before any sample was excluded. The quality of material was estimated using about 10% of the total samples received (selected randomly throughout the collection) by performing: i) a quality check by migration on a 1% agarose gel to ensure the samples were not degraded, ii) a standard PCR amplification reaction on the samples to ensure that the genomic DNA was free of PCR inhibitors, iii) a PCR test to verify the gender of the individual (63). All samples with concentrations below 20 ng/µL, or a major quality problem (degradation and/or amplification problems) were systematically excluded from the study.

After quality control, DNA samples were aliquoted in 96-well plates (JANUS liquid handling robot, Perkin Elmer) for genotyping; sample tracking was ensured by a systematic barcode scanning for each sample. Two DNA positive controls were systematically inserted in a random fashion into the plates. Genotyping was performed on Illumina HumanOmni5-4v1 chips, on a high throughput Illumina automated platform, in accordance with the standard automated protocol of Illumina ® ‘Infinium HD Assay (Illumina ®, San Diego, USA).The PCR amplification of the genomic DNA was performed in a dedicated (« pre-PCR ») laboratory. Fragmentation and hybridization of the DNA on the chips were performed in a dedicated (« post-PCR ») laboratory. Several quality controls were systematically included during the process, such as visual inspection of the DNA pellets after precipitation, visual inspection of the deposited cocktail of reagents for hybridization, systematic verification of the temperature of the heating block during the extension and imaging steps. Reading of the chips was performed on iScan+ scanners (Illumina®, San Diego, USA).Primary analysis of the genotyping results was done using the GenomeStudio software (Illumina®, San Diego, USA). The analysis of the internal controls provided by Illumina and the randomly distributed positive controls allowed the validation of the technological process.

Standard control steps and criteria (64) in GWAS were then applied to data. For DNA samples, individuals with discordant genotypic and reported sex were removed (n=5); heterozygosity versus sample call rate were examined, leading to removal of one clear outlier; all remaining samples had a call rate >0.97 (mean = 0.998) and were kept for analysis. A genetic relationship matrix (GRM) *(65)* was calculated on common variants (MAF > 0.05) and thinned data (r^2^<0.2) to identify duplicates and examine relationships between individuals. A relatively high relatedness was observed in both cohorts which was expected as most of the recruitment was made in rural villages. One pair of individuals was removed because of an unexpectedly high kinship coefficient (Ф = 0.27, corresponding to siblings). All others were kept for association analysis (maximum kinship coefficient Ф = 0.16). Markers with call rate <0.98, MAF <0.01, Hardy-Weinberg equilibrium (HWE) test *P*<10^−8^ (calculated on unrelated individuals) were filtered out, as well as all non-autosomal SNPs. Furthermore, a set of 21 pairs of samples (duplicated DNA samples) were used to identify and remove SNPs with poor reproducibility (4986 SNPs showing at least one discordance). After these quality control steps, the total sample contained 2,363,703 markers for 775 children.

### Population stratification

A principal component analysis (PCA) was performed to evaluate potential population stratification in our samples. The analysis were performed on 624 pairwise unrelated individuals (Ф< 0.05), after filtering on allele frequency (MAF >0.05) and LD thinning (r^2^<0.2). The 151 remaining individuals were then projected onto the principal components.

In order to compare our two Beninese populations with other West African populations, a second PCA was performed including West African populations of the KGP. Genetic data were merged on common positions, filtered and thinned as for the first PCA. Only 100 unrelated individuals from each of the two study cohorts were selected, to give them the same weight as KGP samples in PCA. The remaining individuals were projected on the factorial plan thus obtained.

### Imputation

Imputation of the two cohorts together was performed on the Michigan Imputation Server (66). The reference allele was homogenized to the reference genome by converting the genotypes to the positive strand when necessary; duplicated SNPs, indels, monomorphic SNPs, and SNPs with inconsistent reference allele with the reference genome were removed. The resulting genotype data containing 2,539,302 SNPs were uploaded to the Michigan Imputation Server. A last QC step was performed on the server by comparing allele frequencies observed in the uploaded data and in the African reference panel from 1000 Genomes v5(67), and flipping strand for variants showing significant difference in allele frequency (χ2 statistics greater than 300). After this step, there was a high matched between genotype data and the reference panel (S5 Fig).

Genotype data were finally imputed using the minimac3 algorithm provided by the Michigan Imputation Server. We selected SHAPEIT v2 for prephasing (68,69) and all haplotypes from 1000 Genomes v5 (67) as the reference panel. We filtered out imputed SNPs with a squared correlation (R^2^) between input genotypes and expected continuous dosages below 0.8, or with a MAF below 0.01, which yielded an imputed genotype data set of 15,566,900 variants.

### Adjusting on relevant epidemiological and environmental factors

Association studies were performed with RMM and RMI. A mild malaria attack was defined as a positive RDT or TBS along with fever (axillary temperature ≥ 37.5°C) or a history of fever in the preceding 24 hours. Each mild malaria attack was recorded with its precise date of event and children were considered not to be at risk of malaria within the 14 days after receiving an anti-malarial treatment. Malaria infections included both symptomatic and asymptomatic infections. A positive malaria diagnostic TBS without any fever or history of fever in the preceding 24 hours was considered as an asymptomatic infection if the child did not experience a clinical malaria in the following three days.

The risk of infection greatly varies over the year depending on the season and malaria transmission level. This variation can be accounted for using the time dependent risk of exposure which was defined for each infant in a previous work (Cottrell et al. 2012). To do so, we used a Cox model for recurrent events, with a rate of events at time *t* (Therneau et Grambsch 2000)

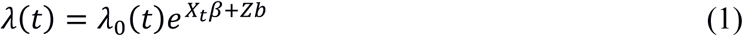

where *X*_*t*_ is a design matric for (possibly time dependent) covariates with a fixed effect, *Z* is a matrix of indicator variables designed for including a vector of random individual effects *b* with variance proportional to the GRM 2*Φ*, allowing to account for cryptic relatedness and population structure. The matrix *X*_*t*_ includes covariates (sex, birthweights, risk of exposure at time *t*, etc). This model has the advantage of both taking into account incomplete follow-up and all the malaria infections, while incorporating time-dependent susceptibility variables.

To fit this model, the follow-up was divided in month intervals following scheduled visit. For each event we considered the risk of exposure estimated at the previous scheduled visit. A stepwise backward strategy was used to select relevant covariates. Factors considered were individual factors (sex, birth weight), health center, factors related to malaria during pregnancy (placental malaria, infection during pregnancy - replication cohort only-, intake of intermittent preventive treatment – discovery cohort only-, arm of clinical trials - replication cohort only-), parity (primigravida vs multigravida), maternal anemia at delivery, use of bednet during the follow-up (categorical variable based on mother declaration during the follow-up), risk of exposure and transmission season (a time-dependent variable in four categories, both follow-up spanning over 3 years: dry season, rainy season 2007, rainy season 2009 and rainy season 2010 for example for the discovery cohort) and socio-economic factors (mother educational level, marital status). The final model included only covariates significant at *p* ≤ 0.05).

### Genome-wide association analyses

It would have been appealing to use the Cox model (1) for the genome-wide analysis, including the genotype of each SNP to be tested for association in the covariates matrix *X*_*t*_. However the computational burden inherent to this modeling (with a non-sparse GRM 2*Φ*) makes it inappropriate for the large number of tests in GWAS.

We thus used the Best Linear Unbiased Predictor (BLUP) 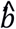 of *b* obtained from the fitted model (1), including all covariates selected by the stepwise backward procedure. This value 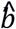 is an “individual frailty”, corresponding to individual effects which are not explained by the epidemiological covariates. To investigate genetic factors involved in these individual effects, they were analyzed using a linear mixed model 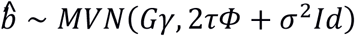 where *G* is the vector of genotypes at the SNP to be tested, coded as 0, 1 or 2 according the number of alternate alleles, which corresponds to an additive model on the log hazard-ratio scale.

This analysis was performed for each genotyped SNP with a MAF > 0.01, as well as with dosages of imputed SNPs. The GRM matrix 2*Φ* used was computed as described in the quality control section. The quantile-quantile plot of the *p*-values was visually inspected and the genomic inflation factor λ (ratio of the median of the observed distribution of the test statistic to the expected median) was calculated to verify the absence of inflation of test statistics due to relatedness and population structure.

SNPs that showed an association at *p* ≤ 10^−5^ in the discovery cohort were selected for replication where the same two steps approach was applied. Gene-based annotations given by ANNOVAR (71) and CADD score (25) were added for these variants in supplementary tables. All highly suggestive signals (around the significant threshold of *p*≤ 5*x*10^−8^ in the discovery cohort, or which replicated at *p* ≤ 0.05) were re-analysed with the mixed Cox model of equation (1) above. For these signals we additionally performed a meta-analysis combining the results obtained by mixed Cox models, using classical method implemented in METAL software *(72)*.

All statistical analyses were done using R software (73), with the package gaston (74) for data quality control and linear mixed models, and the package coxme (23) for the random effects Cox model. Manhattan plots were obtained with the package qqman (75), and regional linkage disequilibrium plot for association signals with Locuszoom Standalone v1.4 (76).

### Gene mapping and biological prioritization

Loci that showed evidence of association at *p* ≤ 10^−5^ with one of the two phenotypes in the discovery cohort, were mapped to gene using the FUMA web plateform (FUMAGWAS v1.3.3; http://fuma.ctglab.nl/) (27), designed for post-processing of GWAS results and prioritizing of genes. FUMA annotates candidate SNPs in genomic risk loci and subsequently mappes them to prioritized genes based on (i) physical position mapping on the genome, (ii) expression quantitative trait loci (eQTL) mapping and (iii) 3D chromatin interactions (chromatin interaction mapping). It incorporates the most recent bioinformatics databases, such as Combined Annotation Dependent Depletion score (CADD score) (25,26), regulatory elements in the intergenic region of the genome (RegulomeDB) (28), Genotype-Tissue Expression (GTEx) and 3D chromatin interactions from HI-C experiments (29).

We considered as genomic risk loci, loci with evidence of replication (table 1) and regions with at least 3 SNPs below the *p*-value threshold of 10^−5^. In the initial step, SNPs in LD (r^2^>0.01, estimated from AFR reference panel of KGP phase 3) in a 250 Kb windows were considered as belonging to the same risk locus. Thereafter, a set of candidate SNPs were selected, based on LD (r^2^ >0.6) with the lead SNP (defined as SNP with the lowest p-value) or with replicating SNPs from our dataset and from AFR population data of KGP phase 3.

Positional mapping was performed using maximum distance from SNPs to gene of 10 kb. Criteria to define deleterious SNPs was a CADD score >12.37 which indicates potential pathogenicity or a RegulomeDB score ≤ 2 which indicates that the SNP likely lies in a functional location (categories 1 and 2 of Regulome classification identified as “likely to affect binding”). EQTL mapping was performed using eQTL data from tissue types relevant for non-severe malaria. Search for eQTL associations were carried out in Blood eQTL (77), BIOS QTL (30), eQTLGen, three databases of e-QTL associations identified in blood, and in the following tissue types of GTEx v7 database: whole blood, cells-transformed fibroblast, cells-EBV-transformed lymphocytes, liver, skin (sun exposed or not) and spleen. Chromatin interaction mapping was performed from Hi-C experiments data of five tissues/cell types identified as relevant for non-severe malaria: liver, spleen GM12878, IMR90 and trophoblast-like-cell. Hi-C experiments identified chromatin interaction between small chromosomal regions two by two. In this analysis a candidate SNP is mapped to a gene if a significant interaction is observed between a first region containing the candidate SNP and the second one where the gene is located. FDR was used to correct for multiple testing (FDR <0.05 for eQTL association and FDR <1 × 10^−6^ for chromatin interaction, the default values in FUMA).

### Candidate associations

We report associations observed in the discovery cohort for genes (± 200 kb 3’ and 5’) showing the greatest evidence of association with malaria in the literature. Genes previously associated with mild malaria phenotype were selected from the review by Marquet (7): *HbS* and *HbC* alleles of *HBB, IL3* from the 5q31-q33 region, *TNF, NCR3* from 6p21-p23 region, and *IL10*. Association was not assessed for the *HP* gene and apha-thalasemia condition, because the structural variants involved are not directly accessible through high density genotyping array (78). We also checked significance for genes previously associated with severe malaria that were confirmed by recent GWA and multi-center studies: *ABO, ATP2B4*, and *FREM3/GYP* locus (17–19,22).

## Acknowledgements

This research is a collaboration between the CEA/ Jacob/CNRGH and the IRD/UMR216. We wish to thanks the collaborators of the CERPAGE who participated actively in the longitudinal follow-ups. Longitudinal follow-ups were funded by the ANR grants (ANR-SEST 2006 040-01 and ANR-PRSP 2010 012-001); the Ministère de la Recherche et des Technologies, France (REFS Nu2006-22) and the IRD. We made use of data previously generated in the MiPPAD study (EDCTP-IP.07.31080.002). The genome-wide scan was supported by the CNRGH. We are grateful to the Genotoul bioinformatics platform Toulouse Midi-Pyrenees (Bioinfo Genotoul) for providing computing and storage resources.

