## Supplemental Data for "First genome-wide association study of non-severe malaria in two birth cohorts in Benin"

### Supplementary data

**Table S1 - Description of main characteristics of Tori-Bossito cohort (572 singletons followed > 28 days) for infants included or not in the GWAS**

| Characteristics | GWA sample (n=525) | Infants excluded from analysis (n=47) | P-value |
|---|---|---|---|
| **Health center n(%)** | | | |
| Tori Avamè | 175(33.3) | 13(27.7) | |
| Tori Cada | 256(48.8) | 20(42.6) | |
| Tori Gare | 94(17.9) | 14(29.8) | 0.1358 |
| **Ethnic group[a] n(%)** | | | |
| Tori | 380(73.6) | 28(59.6) | |
| Fon | 55(10.7) | 5(10.6) | |
| Others | 81(15.7) | 14(29.8) | 0.04406 |
| **Maternal characteristics** | | | |
| **Age mean(SD)** | 27.4(5.5) | 27.7(5.9) | 0.73 |
| **Primigravidae n(%)** | 81(15.4) | 5(10.6) | 0.5 |
| **Placental malaria n(%)** | 58(11.1) | 4(8.9) | 0.83 |
| **Anaemia at delivery (Hb < 100g/L) n(%)** | 83(16.0) | 13(28.3) | 0.06 |
| **IPTp[b] use n(%)** | 438(83.4) | 41(87.2) | 0.68 |
| **Education of woman n(%)** | | | |
| No education | 445(84.8) | 37(78.7) | |
| Partial primary | 56(10.7) | 6(12.8) | 0.3664 |
| Complete primary or more | 24(0.05) | 4(8.6) | |
| **Infant characteristics** | | | |
| **Gender male n(%)** | 261(49.7) | 28(59.6) | 0.25 |
| **Low birth weight (< 2500 g) n(%)** | 50(9.5) | 4(8.5) | 1 |
| **Bed net use n(%)** | | | |
| seldom | 36(7.8) | 2(7.4) | |
| freq - | 84(18.2) | 6(22.2) | |
| freq + | 129(28.0) | 7(25.9) | |
| always | 212(46.0) | 12(44.4) | |
| non available | 64 | 20 | |

[a]Ethnic group declared by the mother

[b]IPTp Intermittent preventive treatment during pregnancy with sulfadoxine pyrimethamine.

**Table S2 – Description of main characteristics of Allada cohort (400 singletons) for infants included or not in the GWAS**

| Characteristics | GWA sample (n=250) | Infants excluded from analysis (n=150) | P-value |
|---|---|---|---|
| **Health center n(%)** | | | |
| Attogon | 61(24.4) | 35(23.3) | 0.9038 |
| Sekou | 189(75.6) | 115(76.6) | |
| **Ethnic group[a] n(%)** | | | 0.6837 |
| Aïzo | 170(68.0) | 108(72.0) | |
| Fon | 55(22.0) | 28(18.7) | |
| Others | 25(10.0) | 14(9.3) | |
| **Maternal characteristics** | | | |
| **Age mean(SD)** | 26.2(5.5) | 25.4(5.4) | 0.13 |
| **Primigravidae n(%)** | 36(14.4) | 27(18.0) | 0.42 |
| **Placental malaria n(%)** | 25(10.1) | 18(12.1) | 0.66 |
| **Anaemia at delivery (Hb < 100g/L) n(%)** | 40(16.0) | 31(20.7) | 0.29 |
| **IPTp[b]group(%)** | | | 0.1262 |
| SP | 77(30.8) | 61(40.7) | |
| MQ | 173(70.2) | 89(60.3) | |
| **Education of woman n(%)** | | | 0.06 |
| No education | 168(67.2) | 115(76.7) | |
| Primary or more | 82(32.8) | 35(23.3) | |
| **Marital status** | | | 0.7333 |
| married | 245(98.0) | 146(97.3) | |
| celibate | 5(2.0) | 4(2.7) | |
| **Indicators of socioeconomic status:** | | | |
| having electricity n(%) | 87(34.8) | 51(34.0) | 0.9567 |
| having a television n(%) | 80(32.0) | 53(35.3) | 0.565 |
| having a refrigerator n(%) | 13(5.2) | 3(0.2) | 0.1876 |
| rewarding activity of woman n(%) | 113(45.2) | 64(42.7) | 0.6966 |
| **Infant characteristics** | | | |
| **Gender male n(%)** | 121(48.4) | 68(45.6) | 0.67 |
| **Low birth weight (< 2500 g) n(%)** | 15(6.0) | 21(14.0) | 0.01 |
| **Bed net use n(%)** | | | |
| seldom | 11(4.4) | 3(5.4) | |
| freq - | 29(11.6) | 3(5.4) | |
| freq + | 29(11.6) | 10(17.9) | |
| always | 181(72.4) | 40(71.4) | |
| non available | 0 | 39 | |

[a] Ethnic group declared by the mother

[b] IPTp Intermittent preventive treatment during pregnancy in MiPPAD trials. Women were randomised to receive two doses of IPTp: SP, sulfadoxine-pyrimethamine; MQ, mefloquine.

**Table S3 - Random effects Cox models used to adjust for covariates, discovery cohort.**

| Covariates | Mild malaria attacks | | Malaria infections | |
|---|---|---|---|---|
| | HR (95%CI) | *P* | HR (95%CI) | *P* |
| **Health center** | | | | |
| Tori Avamè | Reference | | Reference | |
| Tori Cada | 2.38 (1.90-2.97) | 1.4e-14 | 2.55 (2.03-3.19) | 1.1e-16 |
| Tori Gare | 1.52 (1.14-2.03) | 4.3e-03 | 1.77 (1.32-2.35) | 9.0e-05 |
| **Risk of exposure** | 1.26 (1.16-1.37) | 6.4e-09 | 1.24 | 1.4e-09 |
| **Transmission season** | | | | |
| dry season | Reference | | Reference | |
| rainy season 2007 | 1.76 (1.06-2.90) | 2.8e-02 | 2.15 (1.38-3.36) | 6.8e-04 |
| rainy season 2008 | 1.46 (1.15-1.87) | 1.7e-03 | 1.62 (1.31-2.00) | 6.2e-06 |
| rainy season 2009 | 3.27 (2.48-4.31) | 1.1e-16 | 2.57 (2.00-3.28) | 7.9e-14 |

HR, Hazard Ratio. Final model included covariates associated at P<0.05. Covariates selection lead to the same covariates set for both traits. Models were performed on 554 children followed more than 3 months, for whom 1072 malaria infections, among them 811 mild malaria attacks were observed.

**Table S4 - Random effects Cox models used to adjust for covariates, replication cohort.**

| Covariates | Mild malaria attacks | | Malaria infections | |
|---|---|---|---|---|
| | HR (95%CI) | *P* | HR (95%CI) | *P* |
| **Health center** | | | | |
| Attogon | Reference | | Reference | |
| Sekou | 1.51 (1.16-1.96) | 1.6e-03 | 1.33 (1.02-1.72) | 0.02 |
| **Risk of exposure** | 1.50 (1.28-1.75) | 2.9e-07 | 1.50 (1.29-1.74) | 3.9e-08 |
| **Transmission season** | | | | |
| dry season | Reference | | Reference | |
| rainy season 2011 | 1.11 (0.85-1.43) | 0.44 | 1.19 (0.94-1.50) | 0.13 |
| rainy season 2012 | 1.53 (1.23-1.91) | 1.1e-04 | 1.41 (1.16-1.72) | 5.5e-04 |
| **Marital status** | | | | |
| Celibate | Reference | | Reference | |
| married | 0.47 (0.26-0.86) | 0.01 | 0.49 (0.25-0.96) | 0.03 |
| **Education of woman** | | | | |
| No education | Reference | | Reference | |
| Primary or more | 0.74 (0.59-0.94) | 0.01 | 0.78 (0.62-0.99) | 0.04 |

HR, Hazard Ratio. Final model included covariates associated at P<0.05. Covariates selection lead to the same covariates set for both traits. As few events were recorded in rainy season 2013 (end of the follow-up at the beginning of this rainy season) they were grouped with rainy season 2012. Models were performed on 295 children followed more than 3 months, for whom 901 malaria infections among them 687 mild malaria attacks were observed.

**Figure S1. Principal component analysis on study data.**

PCA were performed using 624 unrelated individuals for calculation of principal components (PC) with projection onto of remaining individuals: a) and b) plots of the first four PC by cohort; c) and d) plots of the first two PC by ethnic groups and health centers.

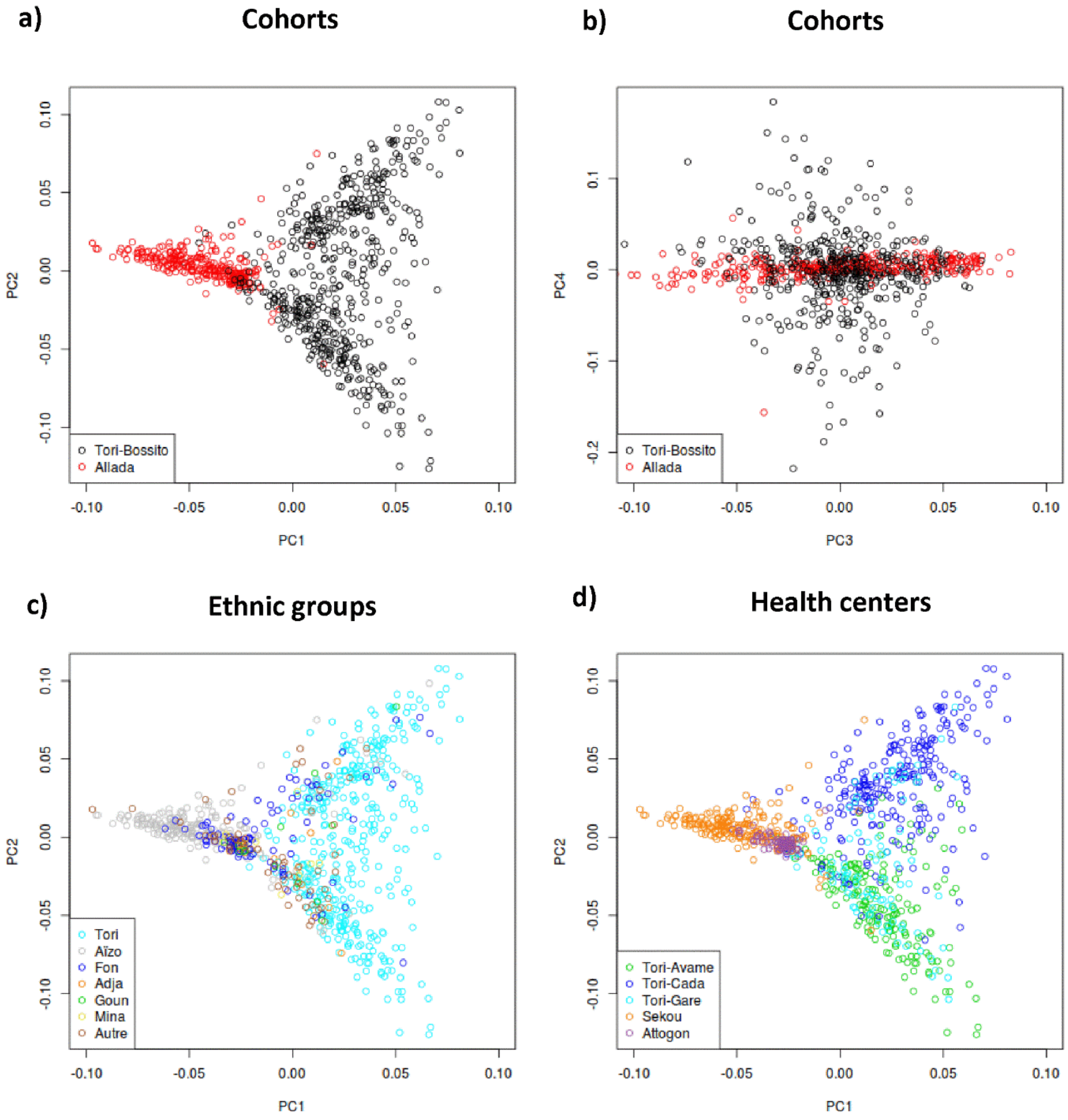

**Figure S2. Principal component analysis including KGP African population.** GWD = Gambian, MSL = Mende in Sierra Leone, LWK = Luhya in Kenya, ESN = Esan in Nigeria, YRI = Yoruba in Nigeria.

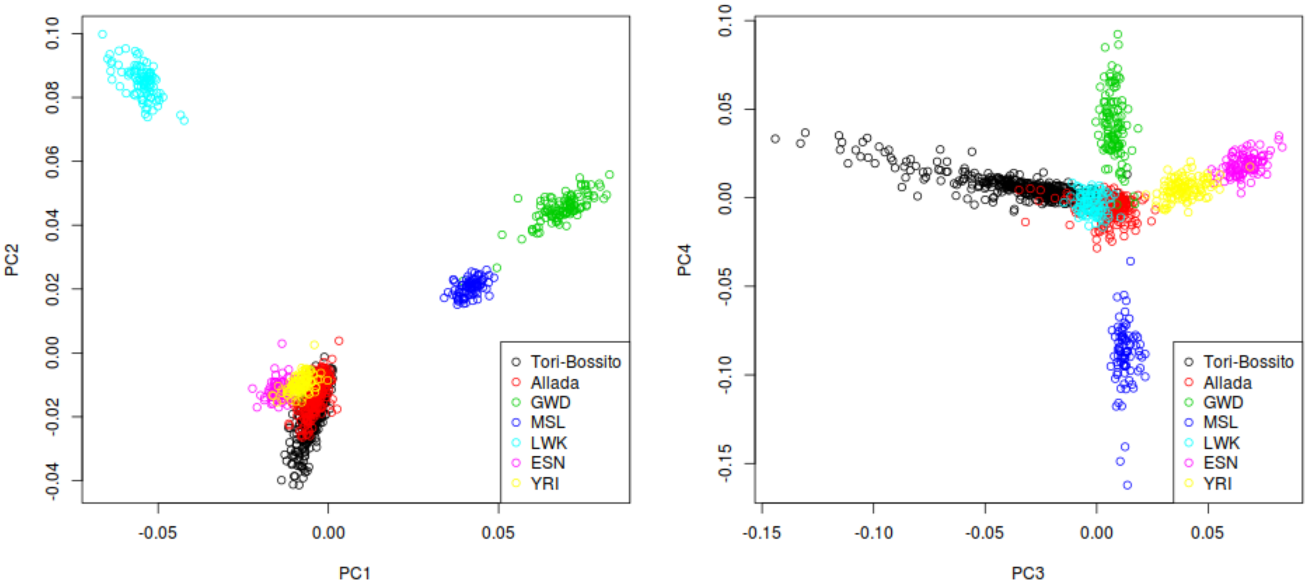

**Figure S3. Annotated regional association plots for main association signals.** Plots show LD calculated on Nigerian Population of KGP dataset (ESN and YRI populations)

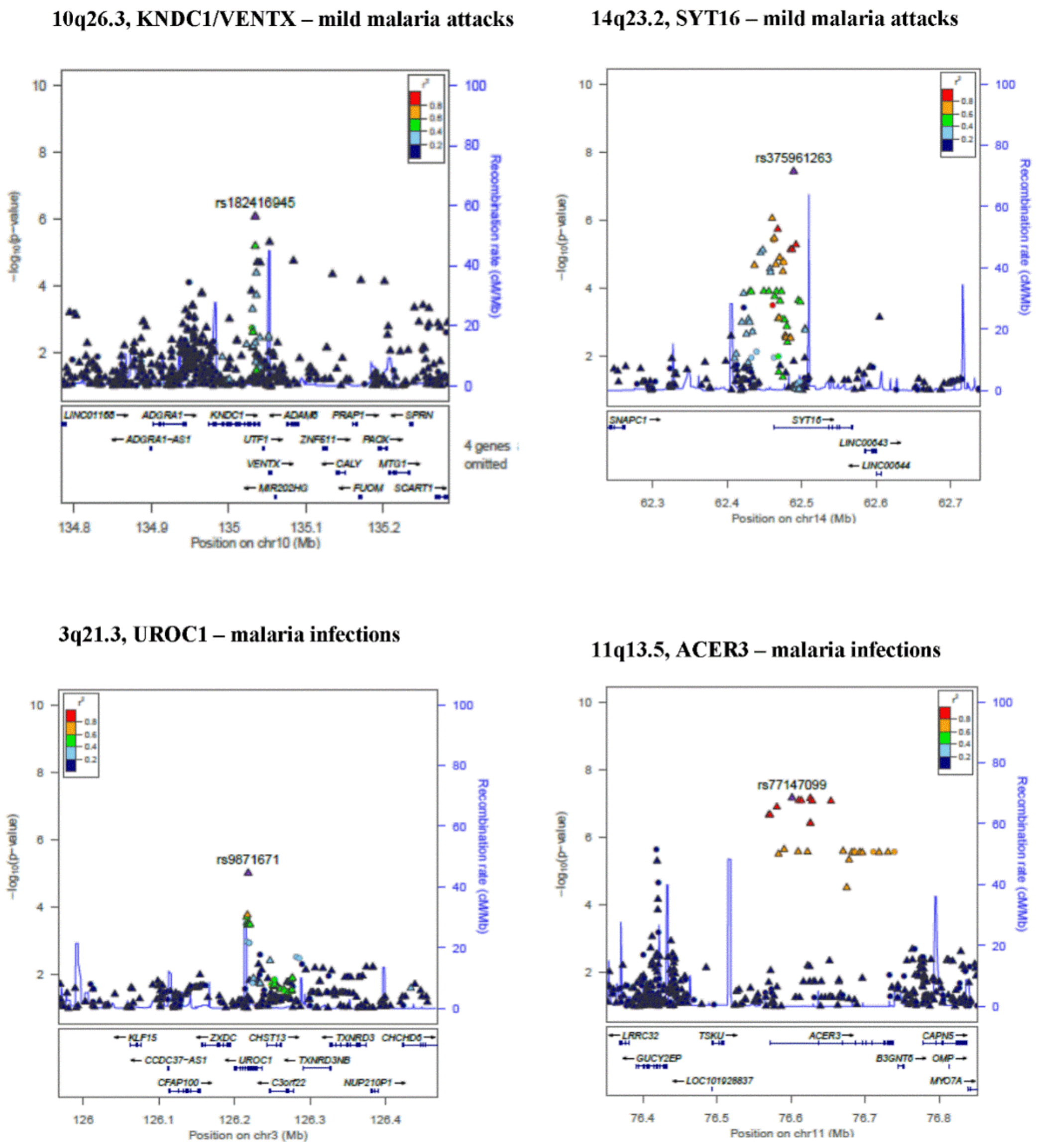

**Figure S4. Results of association analysis after conditioning on the lead SNP, for main association signals.** Plots show LD calculated on Nigerian Population of KGP dataset (ESN and YRI populations)

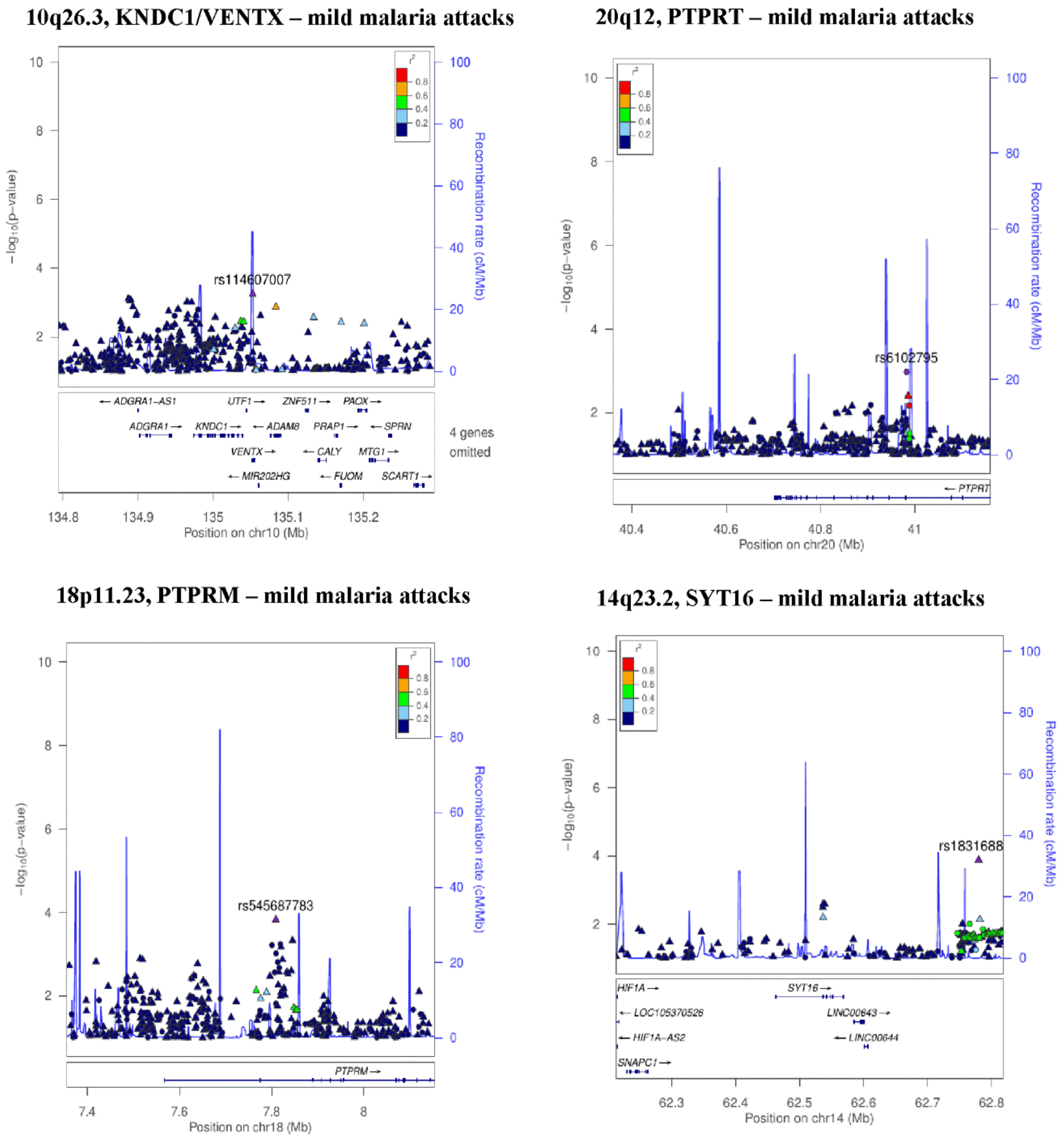

**3q21.3, UROC1 –malaria infections**

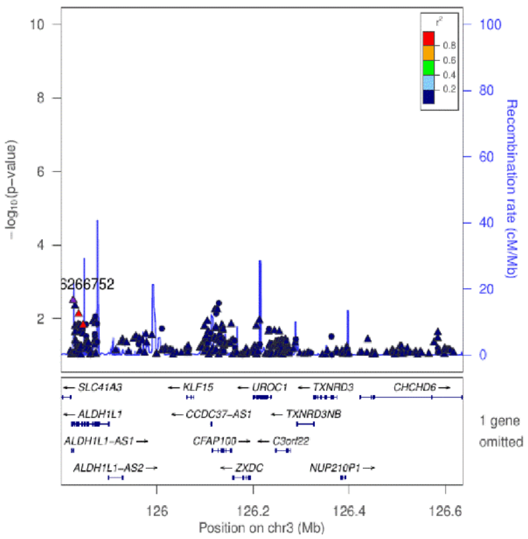


**20q12, PTPRT –malaria infections**

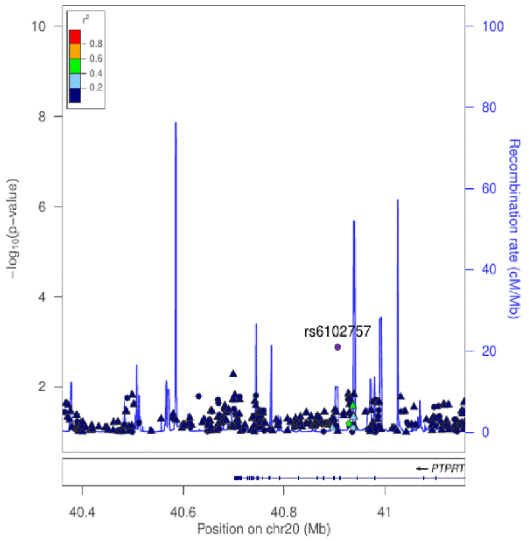


**11q13.5, ACER3 – malaria infections**

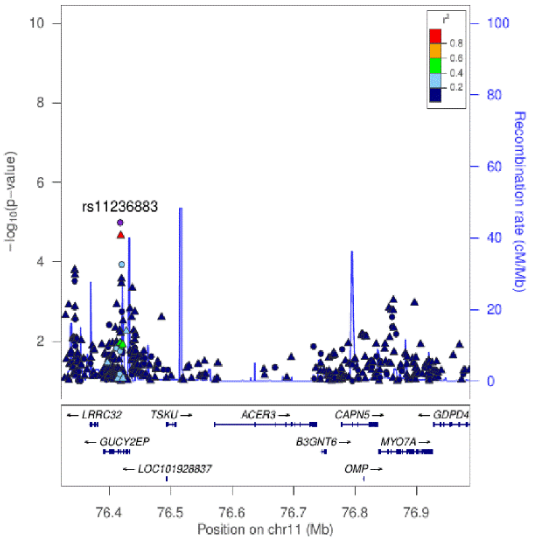


264

**Figure S5. Two cohorts (x-axis) versus African reference panel (y-axis) reference allele frequencies after pre-imputation QC steps.**

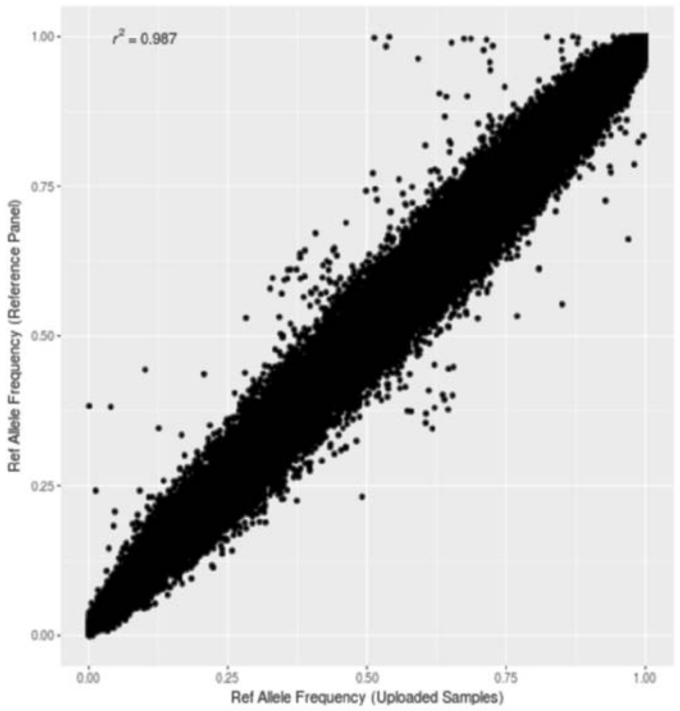
